# Aneuploidy profiling among different types of brain metastases reveals a colon cancer-specific chromosomal signature

**DOI:** 10.64898/2026.10.28.762623

**Authors:** Aparajita Khan, Mengrui Zhang, Susan M. Grimes, Billy T. Lau, Anuja Sathe, Summer Han, Hanlee P. Ji

**Author notes:** These authors contributed equally to this work (Aparajita Khan, Mengrui Zhang).

## Abstract

Brain metastasis is a significant cause of patient morbidity and mortality. There is increasing evidence that genomic instability features such as aneuploidy are associated with metastases to the brain and other organs. We analyzed the genomic instability of brain metastases among different cancer types. This included brain metastases from non-small cell lung, breast, melanoma and colorectal cancer. Our study identified a significantly higher degree of genomic instability in colorectal brain metastases compared to the other types of brain metastases. Nex, we examined the brain metastases for the types of aneuploidy events, namely chromosomal arm imbalances that involved gains or losses. Colorectal cancer brain metastases had specific chromosomal arm imbalance profiles that included gains of 13q and 20q which were significantly more frequent compared to the other cancer types. We examined other colorectal cancer metastatic sites in the lung, liver, lymph nodes and the primary colon. Examining colorectal cancer metastases more closely, aneuploidy gains across either arms for chromosomes 7 and 16 were also higher among brain metastases compared to liver, lung and lymph node metastases. Overall, this study shows that colorectal cancer brain metastasis has specific aneuploidy profiles different than other cancer types as well as being different among other metastatic organ sites.

## INTRODUCTION

An estimated 20% of all cancer patients will develop brain metastases (**BMets**) which leads to substantial morbidity and high mortality [1]. Treatments include surgery, radiation, targeted therapy and immunotherapy. Oftentimes, these treatments provide only limited local control of the metastases. As a result, the median survival is typically months [2]. Even with recent improvements in systemic therapies for treating metastases in other organs, the diagnosis of BMets remain a life-threatening event. Therefore, it is a priority to identify the molecular and genetic features of brain metastasis and improve our understanding of the underlying mechanisms.

The brain is a restrictive, hostile environment for epithelial tumor cell growth [1]. Epithelial tumor cells must overcome multiple biological hurdles representing critical bottlenecks for metastasis [1]. Steps in this process include: (1) vascular transit to the brain while overcoming shear stresses; (2) traversing the blood–brain barrier; (3) adapting to the brain microenvironment which has metabolic constraints, limited immune surveillance and foreign cellular microenvironment. Experimental in vivo selection studies have demonstrated that tumor cells must have specific properties to establish brain metastases [3].

Specific cancers have different rates of brain metastases with non-small cell lung cancer (**NSCLC**), breast cancer (**BRCA**) and melanoma (**MELAN**) having the highest frequency [4–6]. Digestive tract cancers generally have much lower frequencies of BMets. Among these digestive malignancies, colorectal cancer (**CRC**) has the highest rate of BMets albeit much lower than what is observed among NSCLC, BRCA or MELAN [6]. Some of these differences in rates for BMets may be related to a specific genomic features to a given cancer type and their metastatic potential to spread to the brain.

Many cancers have an extensive degree of underlying genomic heterogeneity. This is a result of chromosomal instability (**CIN**) which is a hallmark of cancers, characterized by large chromosomal alterations, often associated with poor prognosis and contributes to metastasis [7]. Among these chromosomal instability features, aneuploidy is prominent, where cells have an abnormal chromosome composition that include increased chromosome number and chromosomal arm imbalances, is a feature of cancer development and progression and associated with clinical outcomes. Up to 90% of solid tumors have aneuploidy, a state where there are dramatic changes in the number of chromosomes as well as gain and losses in smaller segments such as their chromosomal arms [8, 9]. One study showed that different cancer types have unique aneuploidy profiles [10]. Metastatic cancers show higher levels of events aneuploidy compared to primary tumors, indicating that highly aneuploid clones are selected during the metastatic process [10]. In terms of brain metastasis, another study of matched primary tumors and their brain metastases showed evidence of specific chromosome imbalances [11]. Overall, there is evidence pointing to the possibility of recurrent patterns of aneuploidy associated with brain metastasis [12].

We conducted a study comparing the genomic instability and aneuploidy features of BMets. Our study focused on those cancers with a high rate of BMets such as non-small cell lung cancer, breast cancer, melanoma versus a digestive tract cancer, colorectal cancer which has much lower rates of BMets. We identified specific genomic features of BMets among the different cancer types. Our study evaluated different genomic metrics that included the fraction of genome altered (**FGA**), tumor mutational burden (**TMB**), microsatellite instability (**MSI**) and other characteristics among different types of brain metastases versus other metastatic sites and primary tumors. Moreover, we quantified and compared the gains versus loss events on chromosomal arms among the different types of brain metastases and other metastatic sites. Given the rarity of CRC BMets and their having the highest level of genomic instability among the four cancer brain metastases, we focused on CRC BMet aneuploidy patterns and specific chromosomal alterations that are associated with brain metastasis. The rarity of CRC BMets provided us with an opportunity to identify these brain-specific aneuploidy events compared to other CRC metastatic organ sites.

## METHODS AND MATERIALS

### Cancer genomic data from brain metastasis and other metastatic sites

We had three data sets to determine the aneuploidy features from metastatic cancers to the brain. Set 1 had the largest number of tumors from 50 different cancer types. This data came from targeted sequencing of cancer that were part of the MSK-IMPACT and AACR Project GENIE studies [13]. For additional analysis of CRC aneuploidy events, we included two other data sets. Set 2 had exome sequencing of metastases that included 16 patients with metastatic CRC [14]. Among these patients, 11 had brain metastases – some patients presented with initial metastases to other locations, such as, lymph nodes (n = 5), liver (n = 2), lungs (n = 2), and various extracranial sites (n = 3). Five patients had only brain metastases with no prior metastases to the lung or liver. For Set 3, we conducted whole genome sequencing on 14 CRC metastases to either the liver or the brain. There were 10 brain metastases, eight of which had a paired primary CRC and its brain metastasis.

### Whole genome sequencing of colorectal brain metastasis

The Institutional Review Board at Stanford University School of Medicine approved the study (Protocol 44036) and informed consent was obtained. Set 3 samples were obtained from the Stanford Cancer Center Tissue Bank. Genomic DNA was extracted with the E.Z.N.A. SQ DNA/RNA Protein Kit (Omega Bio-Tek). DNA was quantified with Life Technologies Qubit. A 10X Genomics Chromium instrument (Pleasanton, CA) was used for sample preparation according to the manufacturer’s protocol [15]. Genomic DNA was processed and barcoded libraries were prepared without size selection. Barcoded sequencing libraries were generated with either the GemCode or Chromium kits per the manufacturer’s protocol. Libraries were run on an Illumina Novaseq. All samples were aligned against the NCBI build 37 human reference genome using bwa version 0.7.5a-r405 [16] with commands aln and sampe. After alignment, duplicates were marked using Picard MarkDuplicates and the GATK variant calling pipeline steps [17] were followed using standard best practices, including IndelRealigner and BaseRecalibrator. Samples were merged at the indel realignment step. After the UnifiedGenotyper was run, VariantRecalibration and ApplyRecalibration were run to produce filtering information. Only SNVs passing these filters and having a total allelic depth of at least 10 were used to generate a VCF file for the Long Ranger Supplementary phasing mode processing described above. Normal tissue and tumor/dysplastic samples from conventional WGS were compared to identify somatic cancer-specific variants. Only somatic variants passing quality filter and with an overall allelic depth >= 10 were included.

### Determining aneuploidy and chromosomal arm copy number

Among the brain metastases, we determined the amplification and deletion events for each chromosomal arm. We aggregated the chromosomal features from the genomic data sets. To ensure consistency in our analysis across different reference genomes, we updated the segmented copy number data to the genomic coordinates of the latest reference GRCh38. This harmonization was performed using the UCSC LiftOver tool [18]. For this analysis, we used the ASCETS tool which provides chromosome arm changes from genome sequencing data [19]. It uses the genomic segment coordinates, the number of targeted regions from a given segment and the log2 copy number ratio (**LCRs**) compared changes compared to a diploid genome. ASCETS also works on whole genome sequencing data as well. The algorithm divides the alternating LCRs into two groups and then calculates the mean difference between these groups – this determines the variance in log2 ratios across the segment, setting a foundation for more accurate noise threshold determination. The mean differences across all segments and samples established a noise threshold, defined as one standard deviation above the overall mean. We used this threshold to identify significant genomic amplifications and deletions. We identified a chromosomal arm imbalance was based on their adjusted log2 ratios with a burden of change (**BOC**) value for each chromosomal arm. This required a minimum of 70% of the arm segment having a consistent copy number change (e.g., only gain / amplification, etc.). The categories for chromosomal arm changes included gains / amplifications (**AMP**), deletions (**DEL**) or diploid-neutral (**NEU**). The chromosome arms that did not meet this criterion were categorized as not called (**NC**). If the BOC falls below a certain threshold (default at 50%) the arm was denoted as having low coverage (**LOWCOV**).

We evaluated whether the proportion of gain / amplifications or deletions in a specific chromosomal arm in a brain metastasis samples were significantly different among the groups of primary tumors and different metastatic sites. To compare the frequency of chromosomal arm-level gains and deletions between CRC brain metastases and other cancer types, we calculated the proportion of samples harboring each alteration for each chromosomal arm. For each cancer type and chromosomal arm, the proportion was defined as the number of samples with an amplification (or deletion) divided by the total number of samples for that cancer type. The proportion changes (**PropChange**) were calculated as the difference between CRC proportions and comparison group proportions that included NSCLC, breast cancer, melanoma and all other cancer types combined together. Statistical significance was assessed using two-sample proportion tests with Bonferroni correction for multiple testing, and false discovery rate (**FDR**) adjustment was applied using the Benjamini-Hochberg method [20].

We determined other quantitative genomic features including the fraction of genome altered (**FGA**) for each sample. This value provides a broad measure of genomic instability, indicating the extent to which the cancer genome copy number has diverged from a normal diploid genome. The calculation involves summing the total length of segments where the absolute value of the log2 copy number ratio exceeded 0.2, indicating significant deviation from normal diploid copy number [21]. This sum is then divided by the total genomic length covered by the sequencing data – this calculation provides the FGA percentage.

Finally, we analyzed and compared cancer mutations across different patient groups using the maftools package [22]. The mafCompare function was used to determine the statistical differences among different groups. This function employs Fisher’s exact test on all genes between the groups to identify differentially mutated genes. It accounts for multiple testing in gene mutation analysis and the p-values were adjusted using FDR correction. The tumor mutation burden (**TMB**) per Mb was quantified by identifying the total number of nonsynonymous somatic mutations within the exome regions covered by the sequencing data. This calculation includes missense, nonsense, and frameshift mutations. The sum of these mutations was then normalized by the total number of bases sequenced, expressed in megabases (**Mb**).

The microsatellite instability (**MSI**) score was obtained using the MSI-Sensor tool – this program determines the level of microsatellite instability from sequencing read data. It compares the length of microsatellite regions between tumor and normal sequencing data. Microsatellite regions include genomic loci with repetitive s di-, tri-, or tetranucleotide repeats – these microsatellites are prone to accumulating replication errors in DNA mismatch repair-deficient cells. MSI-Sensor provides a quantitative MSI score as the proportion of microsatellite sites showing length variations (e.g., insertions or deletions) in the tumor compared to the normal sample. We determined the tumor purity, the proportion of cancer cells within a tumor sample. For Set 1, tumor purity was estimated through pathologic review of each tumor sample [13]. We also evaluated the time since first metastasis; this was a measure of the interval between the diagnosis of the first metastatic event and the date on which the current tumor sample was resected. We defined the metastatic burden which was as the number of metastatic tumors present in a patient at the time that the metastatic biopsy. This clinical data was available for Set 1. The metastatic events were extracted from the pathologic description which included the anatomic location and number and other de-identified clinical metric [13].

## RESULTS

### Overview of the brain metastasis genomic data

As noted, we had three sets of genome sequencing data. Set 1 had deep targeted sequencing from 50 different cancer types [13]. Sets 2 [14] and 3 had either exome or WGS from CRC. First, we conducted an analysis on the Set 1 genomic data coming from the sequencing of primary and metastatic brain tumors from 421 patients with 18 different cancer types (**Figure 1a**). In addition, there were 7,169 patients with metastases to 21 other organ sites. Then we focused on the brain metastases subset and used this data for the remainder of the analysis. From this Set 1, the four cancers with the highest rates of brain metastases included non-small cell lung cancer with the largest proportion at 46.56% (N = 196), followed by melanoma at 14.96% (N = 63), breast cancer at 9.74% (N = 4) and CRC at 5.23% (N = 22).

**Figure 1.**
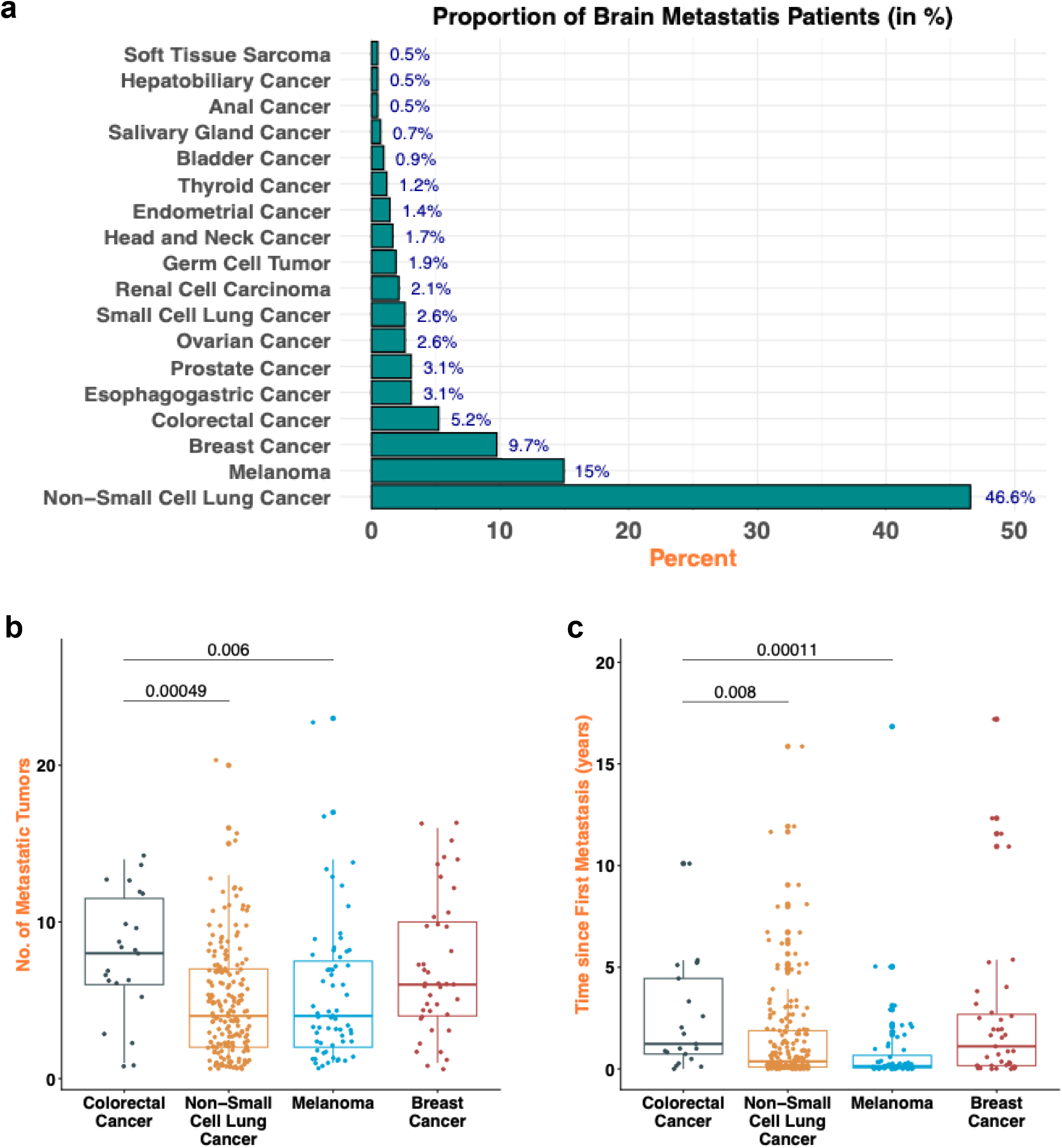
General features of brain metastases. (**a**) Distribution of brain metastasis patients across different primary cancers. The histogram shows that non-small cell lung cancer, melanoma, and breast cancer and colorectal cancer as the top contributors among the others. (**b**) No. of metastatic cancers. The metastatic burden defined as the number of metastatic tumors present in a patient at the time of diagnosis of the brain metastasis. (**c**) Metastatic tumor intervals. This plot represents the time of presentation of the brain metastasis.

From these four cancer types, we determined the number of metastatic sites among different organs other than brain per each patient. For this comparison, we used a total of 4,259 tumors that included CRC, NSCLC, BRCA and MELAN (**Supplementary Table 1**). Among the four different cancer types, 322 were BMets and the remainder were from primary organ site and metastatic sites in organs other than the brain. For this latter group, CRC and breast cancer had the highest number of metastases in other organs (**Figure 1b**). Notably, the average number of metastatic sites among breast cancer patients was comparable to CRC patients. Patients with CRC brain metastases had the highest number of metastatic tumors in other organs, with a mean of 7.95, in contrast to NSCLC (mean of 4.88, adjusted p = 0.001) and melanoma (mean of 5.29, adjusted p = 0.01).

We determined the longitudinal presentation of metastatic sites among patients with brain metastases (**Figure 1c**). For CRC, the interval from the initial metastasis to the detection of brain metastases had an average duration of 2.47 years. This CRC duration was longer than what was observed in NSCLC which had a mean of 1.43 years (adjusted p value = 0.01) and melanoma which had a mean of 0.90 years (adjusted p value < 0.001). Breast cancer had a mean of 2.58 years which did not show a statistical significance difference compared to CRC.

### Genomic features of brain metastases across different cancer types

Next, we focused our analysis on the cancer types with highest number of brain metastases as per Set 1 which had the broadest range of cancer types. For this part of the study, we included the additional CRC data from Sets 2 and 3 for a total of 70 metastatic brain metastases (**Table 1**). Among this group, we compared the fraction of genome altered (**FGA**), tumor mutation burden (**TMB**) and the microsatellite instability (**MSI**) score. CRC brain metastases had a markedly higher FGA (mean [standard deviation - SD]: 0.54 [0.19]) compared to the other cancers including NSCLC (0.41 [0.22], q=0.014), melanoma (0.33 [0.20], q<0.001), and breast cancer (0.42 [0.18], q=0.014) (**Figure 2a**). For TMB, the difference between brain metastases from CRC (8.09 [3.01]) and NSCLC was not statistically significant (13.30 [12.12], q = 0.09). However, melanoma brain metastases exhibited a significantly higher TMB (26.63 [28.70], q<0.001) while breast cancer brain metastases showed a lower TMB (7.32 [7.80], q= 0.019) compared to CRC (**Figure 2b**). This finding is consistent with melanomas having a higher number of mutations as a result of ultraviolet light exposure which is an established mutagenic cancer driver. CRC brain metastases had the highest frequency of MSI, having a significantly elevated mean MSI score of 2.22 when compared to NSCLC (adjusted p = 0.004) and melanoma (adjusted p < 0.001) (**Figure 2c**).

**Figure 2.**
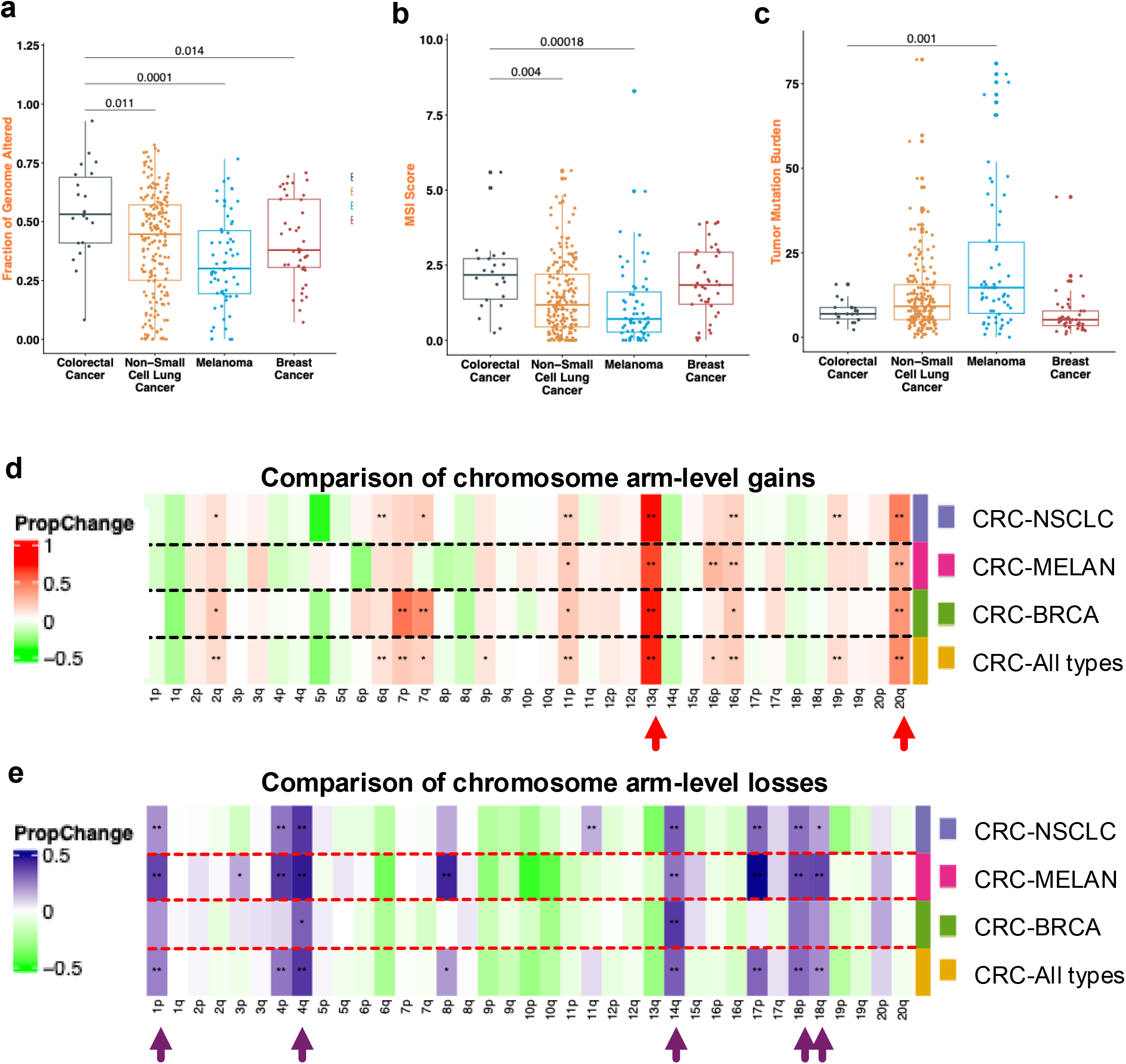
Genomic features of brain metastases. (**a**) Fraction of genome altered (**FGA**). CRC brain metastases exhibit a notably higher FGA compared to lung cancer, melanoma, and breast cancer with a mean FGA of 0.54 for CRC. (**b**) Microsatellite instability (**MSI**) score. CRC brain metastases demonstrate a higher MSI score compared to brain metastases from other primaries, with a mean MSI score of 2.22 for CRC. (**c**) Tumor mutation burden (**TMB**). The analysis reveals a diverse range of mutation burdens, with melanoma exhibiting the highest TMB and breast cancer the lowest compared to CRC. (**d**) Arm level gain comparisons between CRC brain metastases versus other types of cancer with brain metastases. The proportion changes (**PropChange**) were calculated as the difference between the proportions of the CRCs versus the proportions of the other cancer types (CRC-LC, CRC-MELAN, CRC-BRCA, or CRC-ALL) (*P < 0.05, **P < 0.01, FDR-adjusted). The red arrows below the chromosome arm indicate those amplification events which had the highest statistical significance (P < 0.001) and was consistently different across other cancer types. (**e**) Arm-level losses and deletions. The purple arrows below the chromosome arm indicate those deletion events which had the highest statistical significance (P < 0.001).

**Table 1.** Colorectal primary and metastatic cancers.

|  | **Total** | **Set 1** | **Set 2** | **Set 3** |
| --- | --- | --- | --- | --- |
| **Samples** | 1242 | 1111 | 108 | 23 |
| **Patients** | 1141 | 1111 | 16 | 14 |
| **CRC distribution** |  |  |  |  |
| mCRC-Brain | 70 | 22 | 32 | 16 |
| mCRC-Liver | 592 | 584 | 8 | 0 |
| mCRC-Lung | 124 | 124 | 0 | 0 |
| mCRC-Lymph | 79 | 70 | 9 | 0 |
| CRC-Primary | 377 | 311 | 59 | 7 |

### Analysis of aneuploidy between CRC brain metastases and other cancer types

As noted, CRC brain metastases had the highest levels of genomic instability (e.g., FGA) among the different cancer types of brain metastases. This result prompted us to determine if there were specific CRC aneuploidy features compared to the other three cancer types with BMets. We focused on chromosomal arm imbalances, comparing brain metastases from CRC to those from lung (**CRC-NSCLC**), breast (**CRC-BRCA**) or melanoma (**CRC-MELAN**). In addition, we also did a comparison aggregating all the non-CRCs into a single all-inclusive group (**CRC-ALL**) (**Figure 2d**).

We determined the proportion of tumors harboring each chromosomal arm alteration across the genome. For a given cancer type, the proportion metric was defined as the number of tumors with a given chromosomal arm gain or loss divided by the total number of tumors. Next, we determined the difference using the proportional change (**PropChange**) which was the differences for a given chromosomal arm between one type of cancer BMet versus the other. For example, this would involve comparing the arm imbalances for CRC BMets proportions versus the other cancer type’s sample proportions. To determine significance, we applied a two-sample proportion tests with Bonferroni correction for multiple testing and the false discovery rate (**FDR**) adjustment. We identified the chromosomal arm gains and deletions with the highest PropChange value across all three type cancers and identified the subset of events with the highest statistical significance (P < 0.001, FDR-adjusted). We generated heatmaps to visualize the PropChange values across chromosomal arms (**Figure 2d**, **2e**). Generally, we focused on the chromosome arm imbalances with the highest statistical significance (P < 0.001, FDR-adjusted) unless otherwise stated.

### Chromosome arm gains 13q and 20q are significant in CRC brain metastasis

First, we compared the chromosomal arm gains (e.g., amplifications) of the CRC BMets to NSCLC (lung), BRCA (breast) or MELAN (melanoma) (**Figure 2d**, **Supplementary Table 1**). Compared to NSCLC, CRC had five chromosomal arms with significant differential gains. This included chromosome arms 13q and 20q which had the highest proportional change. Next among the events were gains on 16q, 11p and 19p which were statistically significant with more modest levels of enrichment. Compared to BRCA, CRC had four significantly enriched chromosomal arm with significance. This included chromosome arms 13q, 7p and 20q. Compared to MELAN, CRC had four chromosomal arms amplifications. These chromosome arms included 13q, 20q, 16p and 16q. Then, we compared the CRC chromosomal arm gains versus all cancer types. When comparing CRC to the aggregate cohort of all other cancer types (**CRC-ALL**), chromosomal arm gains of 13q, 20q, 16q, 11p, and 19p met the threshold for statistical significance. Overall, chromosome arm gains of 13q and 20q were increased in CRC BMets.

### Comparison of chromosomal arm losses among brain metastases

We examined the chromosomal arm losses / deletions of the CRC BMet profile NSCL, BRCA and MELAN (**Figure 2e**, **Supplementary Table 2**). Compared to NSCLC, CRC BMets had seven significant losses. This included 4q, 14q, 17p, 18p, 4p, 1p and 11q. Compared to BRCA, CRC had only loss of 14q being the only statistically significant deletion event. Compared to MELAN, CRC had nine significant deletions. Prominent copy number losses with higher proportional changes were observed on 17p, 4q, 8p, 4p, 1p, 18q and 18p. Significant deletions were also present on 22q and 14q. Next, we examined the CRC chromosomal arm losses versus all the other cancer types. In the aggregate comparison against all combined non-colorectal cancer types, eight chromosomal deletion events were significantly enriched in CRC compared to all of the others. Consistent with the individual paired comparisons, the most prominent deletions occurred on 4q and 14q. Highly significant, recurrent losses were also observed on 17p, 18p, 4p and 1p. Finally, moderately significant deletions included 18q and 21q.

### Paired co-occurrence of chromosomal arm gains and losses among brain metastases

Among the CRC-BMets compared to the other types of brain metastases we determined the paired combinations of specific aneuploidy patterns involving gains/amplifications or deletions (**Figure 3**). For example, how many brain metastases had imbalances in both chromosome arms 20q and 17p? First, we determined the frequency of pairs of chromosomal imbalances. Ranking was based on the frequency in which both arm events occur in the same sample for a given cancer type. There were 30 aneuploidy imbalance pairs that were present in approximately 40% of the CRC BMets (**Figure 3**). Among the CRC brain metastases, the five highest frequency pairs were as follows: 20q AMP and 17p DEL (59%); 7p AMP and 20q AMP (56%); 13q AMP and 20q AMP (56%); 13q AMP and 17p DEL (54%); 7p AMP and 13q AMP (53%).

**Figure 3.**
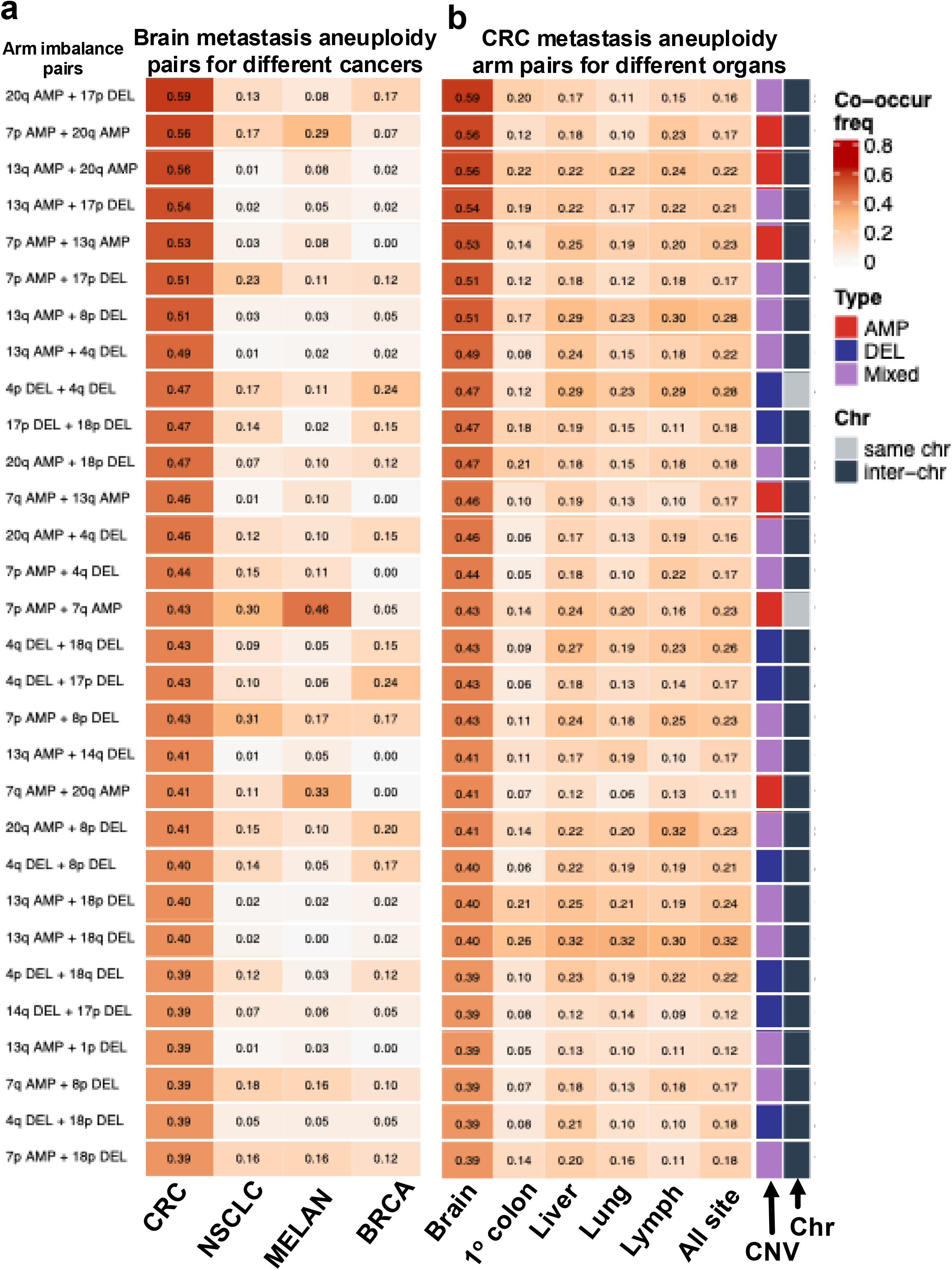
Co-occurrence of chromosomal arm imbalance pairs. Heatmaps show the co-occurrence frequency of the 30 most frequent aneuploidy arm pairs in CRC brain metastases (≥39% of samples). Rows are arm imbalance pairs; color indicates co-occurrence frequency. Annotation bars indicate pair type (AMP, DEL, or mixed) and whether the two arms are on the same chromosome or different chromosomes. (**a**) Frequency of the same pairs among brain metastases from CRC, NSCLC, melanoma (MELAN), and breast cancer (BRCA). (**b**) Frequency of the same pairs among CRC samples from brain, primary colon, liver, lung, lymph node, and all non-brain metastatic sites combined.

Second, we examined the frequency of these CRC BMet aneuploidy arm pairs among NSCLC, MELAN or BRCA BMets (**Figure 3a**). Among the five top frequent pairs seen in CRC BMets, we made a comparison with a binary logistic regression. The event frequency of these five CRC BMet pairs was significantly higher compared to the other cancer BMet types (FDR q < 0.0001). Among the other 25 CRC arm imbalances with a lower frequency, the CRC paired profiles of aneuploidy events were also significantly different (FDR q < 0.001). There were only three exceptions among the other types of BMets. This included MELAN with amplification of chromosome 7 and 7q and 20q arm gain. Also, NSLC had 7p gain and 8p deletion. Overall, these results indicate that CRC BMet had aneuploidy imbalance co-occurring events that distinguished them from other cancer types and their brain metastases.

### Genomic features of CRC brain metastases compared to other sites in different organs

Next, we determined CRC’s genomic features across different metastatic organ sites including the brain, liver, lungs and lymph nodes (**Table 1**, **Figure 4**). The FGA was the highest in CRC brain metastases, with a median of 0.53, compared to the liver (median: 0.22, adjusted p value < 0.001), the lung (median: 0.17, adjusted p value < 0.001), the lymph nodes (median: 0.20, adjusted p value < 0.001) and the colon where the primary tumor originated (median: 0.09, adjusted p value < 0.001) (**Figure 4a**).

**Figure 4.**
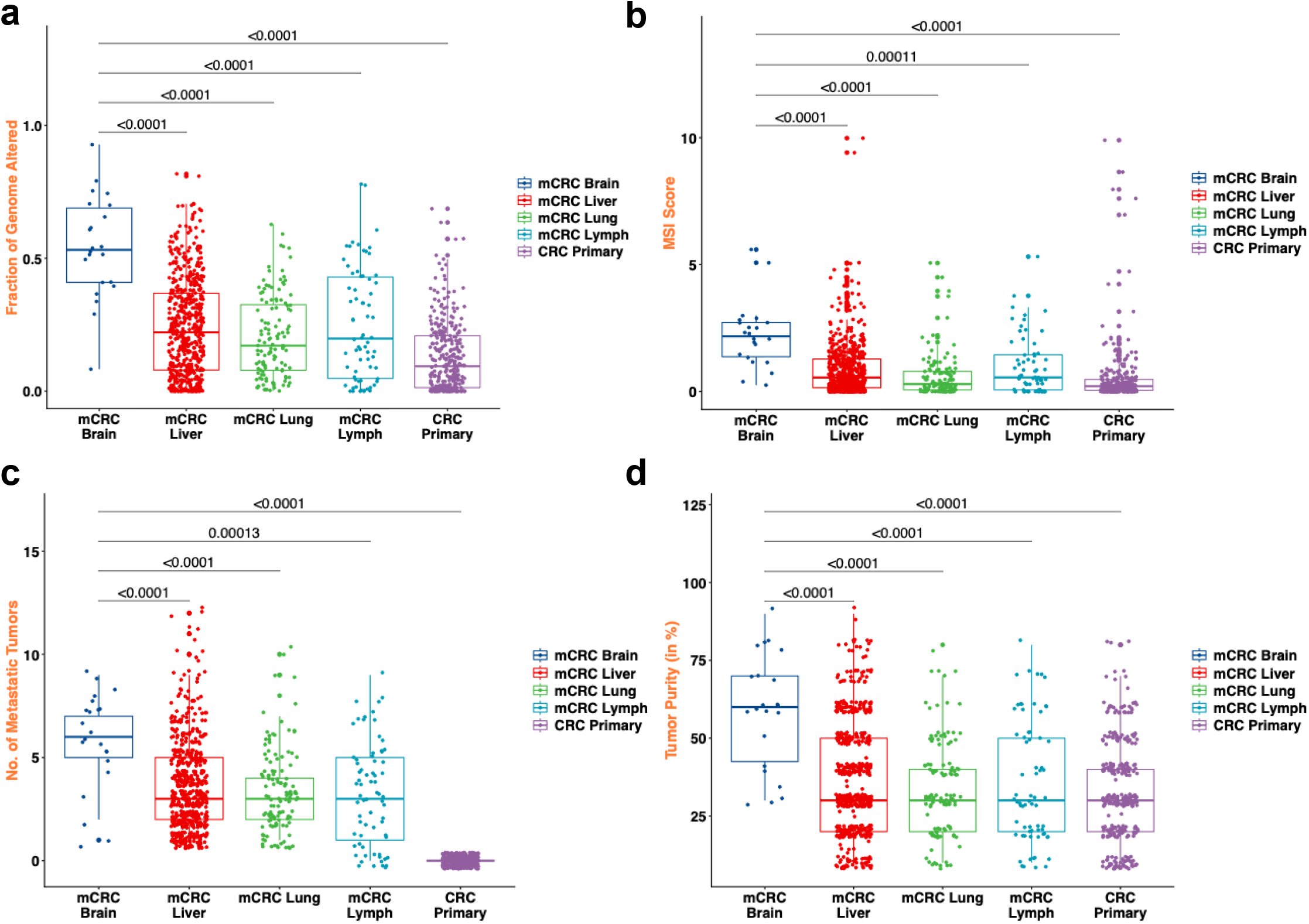
Genomic features of CRC brain metastases compared to other organ sites. (**a**) Fraction of genome altered (**FGA**). Significantly higher FGA was observed in CRC brain metastases compared to other metastatic sites. (**b**) Microsatellite instability (**MSI**) score. The MSI Score varies significantly across different metastatic sites of CRC. Brain metastases exhibit a higher median MSI Score compared to other metastatic sites. (**c**) No. of metastatic tumors: The number of metastatic tumors in CRC brain metastases is significantly higher compared to other metastatic sites. (**d**) Tumor purity. The proportion of cancer cells is significantly higher in CRC brain metastases compared to other metastatic sites.

For those metastatic CRCs with MSI, there was significant variation across the different metastatic locations. Brain metastases had a higher median MSI score of 2.17 compared to the liver (median: 0.57, adjusted p value < 0.001), the lung (median: 0.30, adjusted p value < 0.001), the lymph node metastases (median: 0.72, adjusted p value = 0.003) and the colon origin of the primary tumor (median: 0.28, adjusted p value < 0.001) (**Figure 4b**).

We determined the number of CRC metastatic sites for each patient. For patients with CRC BMets, the median number of other organs with metastatic sites was eight. For those patients without BMets but with metastasis to other organs (e.g., liver, lungs or lymph nodes), the median number of organs with metastases was four. This was a statistically significant difference (adjusted p value < 0.001) (**Figure 4c**) and indicates that patients with BMets have a higher number of metastatic sites in other organs.

Tumor purity was significantly greater in CRC brain metastases, with a median of 60%, in contrast to liver (median: 30%, q < 0.001), lung (median: 30%, q < 0.001), lymph node metastases (median: 30%, q < 0.001) and primary CRC tumors (median: 30%, q < 0.001). This result indicated that CRC brain metastases contain a higher proportion of tumor cell fraction when compared to the other metastatic sites and primary tumors (**Figure 4d**).

### Chromosomal aneuploidy patterns of CRC among different metastatic organ sites

We determined if the CRC BMets had different and specific aneuploidy events compared to primary colon tumors or CRC metastatic tumors in the liver, lymph nodes and the lungs. For this analysis we had a total of 1242 CRC samples with 70 BMets with the remainder coming from primary and metastatic sites (**Table 1**). Statistical significance for the PropChange value was assessed with the same methods used in comparing among different cancer types (**Methods**). Specifically, we identified chromosomal arm gains or losses with the highest PropChange value across all three type cancers and greatest statistical significance (P < 0.001, FDR-adjusted) (**Supplementary Tables 1 - 4**). We generated heatmaps to visualize the statistically significant PropChange among the different chromosomal arms (**Figure 5**).

**Figure 5.**
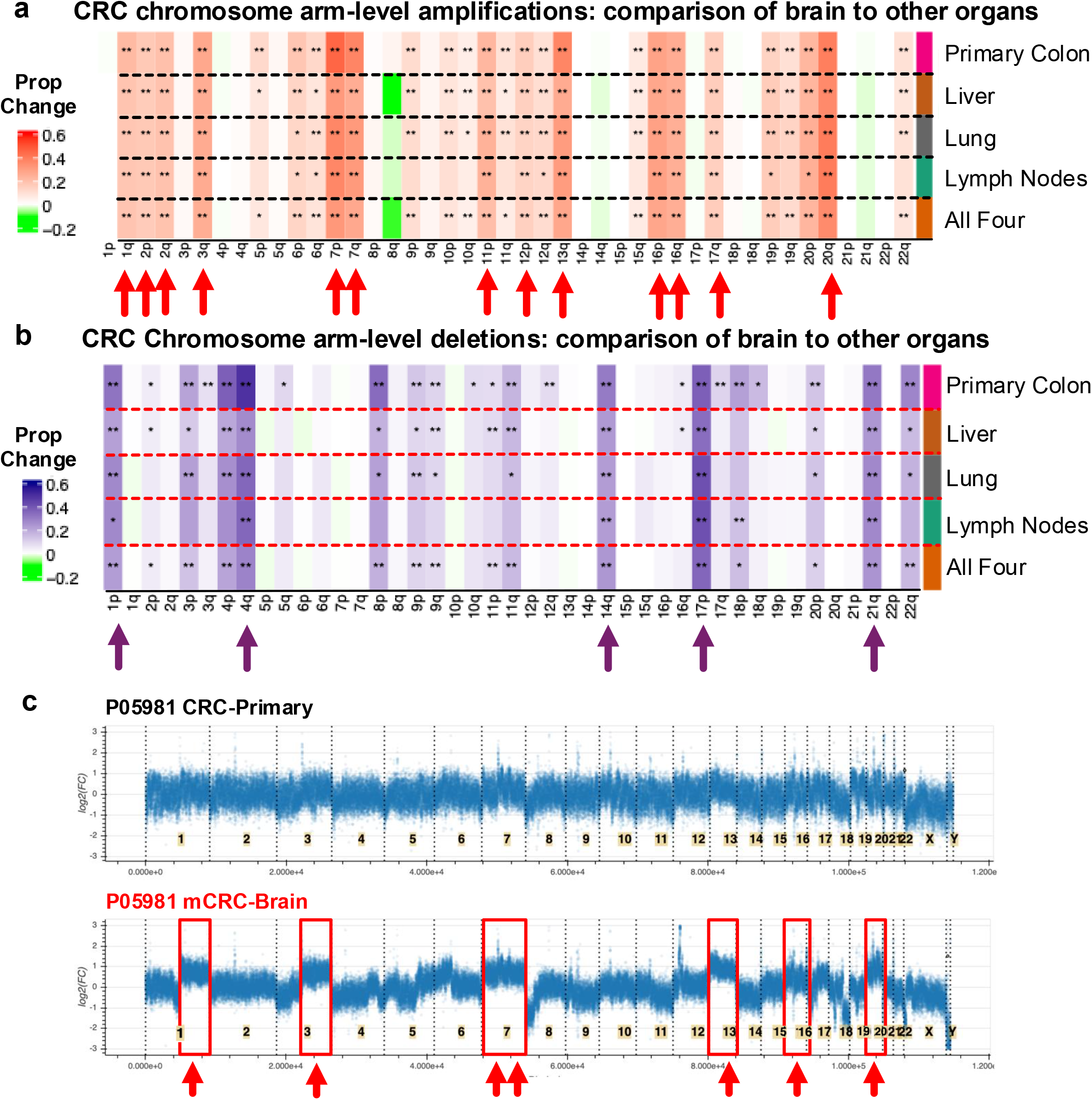
Colorectal cancer aneuploidy feature comparison between brain metastasis and different metastatic organ sites. For each cancer type and chromosomal arm, the proportion was defined as the number of samples with an amplification or deletion divided by the total number of samples for that cancer type. We generated heatmaps to visualize the PropChange values across chromosomal arms, with asterisks indicating statistical significance (*P < 0.05, **P < 0.01, FDR-adjusted). Comparisons between CRC brain metastases versus other CRC metastatic organ sites. Arrows below the chromosome arm indicate those events which had statistical significance (P < 0.001, FDR-adjusted) and was consistently different across other cancer types. The PropChange range is value across 0.6 to -0.2. (**a**) Arm-level gains. (**b**) Arm-level losses. (**c**) Whole genome analysis of the patient-matched primary CRC versus BMet showing these specific aneuploidy features. The BMet had gains in chromosome arms that included: 1q; 3q; 4q; both 7p-q; 13p and q; both 16 p-q; both 20p-q. None of these chromosome alterations were present in the primary CRC. Frequent features from the larger study are highlighted in red boxes and arrows.

We identified the chromosomal arm gains with (1) the highest PropChange value across all three type cancers and (2) statistical significance (P < 0.001, FDR-adjusted). BMets had extensive arm imbalances for nearly all chromosomes (P < 0.001, FDR-adjusted), much more so than the primary CRCs from the colon and the CRC metastases in other organs (**Supplementary Tables 1 and 2**). With these criteria the following chromosomal gains were identified: 1q, both 2p-2q, 3q, both 7p and 7q, 11p, 12p, 13q, chromosome 16 (p and q), 17q and 20q (**Figure 5a**). Among these BMet gains, the top four were 13q (75%, N = 53), 20q (72.9%), 7p (67.1%) and 7q. We also examined the results to determine which chromosome imbalances had the highest frequency in BMets and lower frequency among the other sites (primary colon, liver, lung, lymph nodes). As noted, when making comparisons against other cancer types with brain metastases, gains in 13q and 20q gains were notable findings. Overall, this result points to the aneuploidy gain events being different in brain metastases compared to other sites.

In terms of deletions, the most frequent loss events with the greatest statistical significance included chromosomal arms 1p, 4q, 14q, 17p and 21q among BMets compared to all other sites (primary and other organs) (**Figure 5b**). In contrast to gains, arm losses were not as frequent among BMets (**Supplementary Tables 2 and 4**).

The whole genome analysis of the patient-matched primary CRC versus BMet is an example showing these specific aneuploidy features. We compared the aneuploidy profiles and copy number features from a matched primary colon tumor and BMet, both from the same patient (P05981) (**Figure 5c**). The BMet had gains in chromosome arms that included: 1q; 3q; 4q; both 7p-q; chromosome 13 (p and q); chromosome 16 (p and q); chromosome 20 (p and q). None of these chromosome alterations were present in P05981’s primary CRC. This result suggests that these aneuploidy gains were associated with brain metastases. Notably, gains in 13q and 20q were among the highest frequency events among CRC BMets and had a high co-occurrence.

Finally, we compared the co-occurrence of chromosomal arm losses among the different CRC metastatic sites. As noted previously, we identified the most frequent pairs of aneuploidy events among CRC BMets and compared those to other cancer types (**Figure 3b**). Then we determined the frequency of the 30 most frequent paired events and compared their frequency among the other CRC metastatic sites. All 30 pairs were significantly more frequent in CRC BMets than in primary tumors. Relative to liver or lung metastases, 29 of the 30 pairs were significantly overrepresented, except for concurrent 13q gain and 18q deletion. Compared with lymph node metastases, 19 of 30 pairs remained significantly enriched. Notably, over 50% of CRC BMets had gains in both 13q and 20q. This result suggests that these gains may play a role in brain metastases.

We compared our results to the study of Golas et al., where they examined a variety of CRC molecular features including chromosomal arm imbalances among BMets as well as liver and lung metastases [12]. Lymph node metastases were not included in their study and they did not examine other cancer types with brain metastases. For this comparison, we used our results about BMet-specific chromosomal arm imbalances that were significantly different compared to liver, lung and lymph nodes. Golas et al. reported the CRC aneuploidy events were prominent across liver and lung metastatic sites and BMets. Concordant BMet features between our study and Golas et al. included chromosome arm gains of 7p, 12p, 13q, both 16p-q and 20q.

### Mutation analysis of CRC brain metastases versus other organ sites

Across the different metastatic sites, we determined if there were any differences among potential pathogenic mutations classes including nonsense, missense, in-frame insertions, frame-shift deletions, frame-shift insertions, and in-frame deletions (**Figure 6**). For this analysis, we used the results from Set 1 with its deep targeted sequencing and highest sensitivity for mutations. **Supplementary Table 7** shows the most frequently mutated genes in brain metastases (>10%) compared to primary CRC tumors. Critical driver genes such as *TP53*, *APC* and *KRAS* were frequently mutated in both groups (**Figure 6**). For example, there was a higher frequency of *TP53* showed a higher mutation rate in brain metastases (82%) compared to primary tumors (63%) but there were no statistically significant differences in the mutation rates for most genes when adjusted for multiple comparisons (adjusted p-value: adjusted p value = 1.00). This pattern was consistent across other major genes, such as *APC* (72.8% vs. 74%, adjusted p value = 1.00) and *KRAS* (64% vs. 40%, adjusted p value = 1.00). Overall, there were some trends such as higher *TP53* mutations in BMets but no changes that reach a threshold of significance.

**Figure 6.**
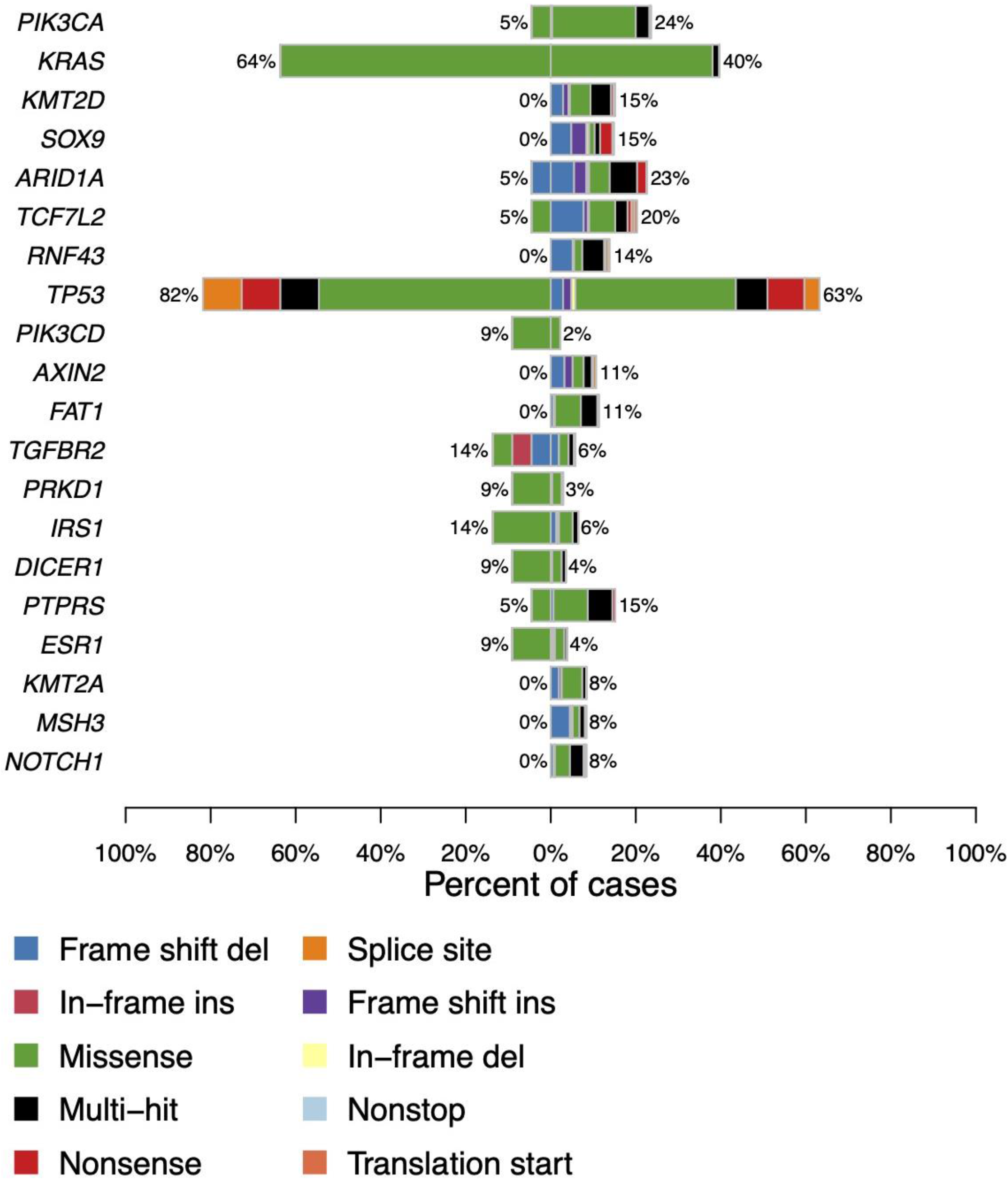
Mutation of brain metastases compared to primary colon cancer. We determined if there were any differences among potential pathogenic mutations classes including nonsense, missense, in-frame insertions, frame-shift deletions, frame-shift insertions, in-frame deletions, splice site mutations and others. Co-barplot of the top differentially mutated genes from Set 1 CRC brain metastases (mCRC-Brain, N = 22; left) versus primary CRC (N = 310; right). Bars show the percent of cases with a mutation; colors indicate variant classification.

Comparing brain to liver metastases (**Supplementary Table 8**), *TP53* mutations were observed at similar rates (82% in brain and liver metastasis, p = 1.00), pointing to no significant difference. Conversely, *KRAS* mutations appeared more frequently in brain metastases (64%) compared to liver metastases (41%), but again, the difference did not achieve statistical significance (adjusted p value = 0.102). Further, both genes *IRS1* and *TGFBR2* displayed differences in mutation rates between the two groups (14% in mCRC-Brain vs. 2% in mCRC-liver, adjusted p value = 0.052), with a p-value slightly above the significance threshold. In the comparison with lung metastases (**Supplementary Table 9**), *TP53* mutations occurred similarly across both sites (82% in brain vs. 83% in lung for *TP53*, adjusted p value =1.00). While *KRAS* was more frequently mutated in brain (64%) compared to lung metastasis (53%, adjusted p value = 0.97), APC showed more frequent mutations in lung metastasis (83% vs. 73% in brain, adjusted p value = 0.97). Similarly, for lymph node metastases (**Supplementary Table 10**), the differences were also not statistically significant, with *TP53* (82% in brain vs. 86% in lymph nodes, adjusted p value = 1) and *APC* (73% in brain vs. 74% in lymph nodes, p=1) showing comparable mutation rates. However, *KRAS* showed a significantly higher mutation rate in brain metastases (64%) compared to lymph node metastases (27%), with an adjusted p-value of 0.016, suggesting a site-specific variance in *KRAS* mutational status between brain and lymph node metastases.

This pattern of non-significant differences in individual gene mutations was consistent across comparisons with liver, lymph node, and lung metastases, suggesting that while large-scale genomic alterations are pronounced, specific gene mutations do not distinctively characterize brain metastases compared to other sites, also underscoring a broader similarity in the mutation landscape despite differences in tumor location and metastatic progression.

## DISCUSSION

We conducted a comparative genomic analysis of brain metastases from different cancer types. This study included lung cancer, breast cancer, and melanoma, which frequently metastasize to the brain, and colorectal cancer in which brain metastases are relatively rare. Our analysis identified distinct aneuploidy events among brain metastases originating from these cancer types. This observation is consistent with recent studies of brain metastases from melanoma and lung cancer, which have also reported increased chromosomal instability in metastatic tumor cells [23–25]. The most striking finding was that CRC BMets exhibited the highest level of genomic instability among all the cancer types examined, as well as distinct chromosome arm-level amplifications and deletions compared with CRC metastases in other organs. These results suggest that extensive aneuploidy may contribute to the metastatic behavior of CRC tumor cells in the brain. These findings are also consistent with a recent independent study [12] and our spatial transcriptomic analysis of CRC-BMets [26], which identified increased chromosomal instability and specific arm-level alterations in metastatic tumor cells.

Aneuploidy is a defining feature of CRC, with recurrent gains and losses of specific chromosomes emerging during tumor development, progression, and metastasis to distinct organ sites [12, 27]. Our results indicate that CRC-BMets have a greater burden of aneuploidy and distinct chromosome-arm alterations even compared with CRC metastases in other organs. In contrast, the somatic mutation landscape of CRC-BMets was largely similar to that of extracranial metastases. Metastasis to the brain is relatively uncommon in CRC and frequently occurs after metastatic disease has developed in other organs [28]. This prolonged disease course may provide opportunities for the selection of tumor subclones with distinct chromosomal configurations. Our results raise the possibility that specific aneuploidy patterns may confer advantages during the biological processes required for brain colonization, including survival during vascular dissemination, extravasation across the blood–brain barrier, and adaptation to the brain microenvironment. Specific chromosome-arm alterations in CRC-BMets could confer metastatic advantages through gene dosage effects. Alternatively, chromosomal instability could influence tumor–microenvironment interactions [29], potentially favoring metastatic growth. Future experimental studies can provide additional insights into the mechanisms by which aneuploidy may contribute to CRC brain metastasis.

Our findings may also have clinical implications. CRC is the third most common malignancy in the United States. While survival rates have generally improved for patients with this cancer, treating those individuals with metastasis remains a daunting challenge. CRC-BMets are relatively rare but are associated with worse outcomes, including increased morbidity and mortality [30, 31]. As treatments for CRC improve and patient survival rates increase, the incidence of BMets is expected to rise [30]. Routine brain MRI screening is not generally recommended for patients with CRC, in part because of the relatively low incidence of brain metastases. Identifying patients at increased risk could therefore help guide surveillance strategies. However, there are currently no established biomarkers for predicting brain metastasis in patients with CRC. Our findings raise the possibility that aneuploidy patterns could help identify patients at increased risk, which should be investigated in future studies using longitudinal genomic analyses of matched primary tumors, extracranial metastases, and brain metastases.

Several limitations should be considered when interpreting our findings. Our study integrates genomic data from multiple cohorts and sequencing platforms, which may introduce differences in the sensitivity and resolution of copy-number detection. Many comparisons involve unmatched tumors, limiting our ability to reconstruct the acquisition of chromosomal alterations during individual patients’ metastatic progression. Clinical differences between patients with brain metastases and those with metastases to other organs, including disease duration, metastatic burden, and treatment history, may also contribute to the observed genomic differences.

## DATA AND CODE AVAILABILITY

Sequencing data has been deposited to dbGAP identifier phs003794. The code and demo data are in the following URL: https://doi.org/10.5281/zenodo.23674551.

## AUTHORS’ CONTRIBUTIONS

AK and MZ was involved in conception and design of the study, development of methodology, acquisition of data, analysis and interpretation of data and writing of the manuscript. AK, MZ, SMG and SA were involved in the acquisition and analysis of data. SH and HPJ oversaw the conception and design of the study, data analysis, interpretation of data and writing of the manuscript.

## ACKNOWLEDGEMENTS

We are grateful to all patients who participated in the study.

## REFERENCES

1. Achrol, A.S., et al., Brain metastases. Nat Rev Dis Primers, 2019. 5(1): p. 5.

2. Deshpande, K., et al., Clinical Perspectives in Brain Metastasis. Cold Spring Harb Perspect Med, 2020. 10(6).

3. Miarka, L. and M. Valiente, Animal models of brain metastasis. Neurooncol Adv, 2021. 3(Suppl 5): p. v144–v156.

4. Owen, S. and L. Souhami, The management of brain metastases in non-small cell lung cancer. Front Oncol, 2014. 4: p. 248.

5. Niikura, N., et al., Brain metastases in breast cancer. Jpn J Clin Oncol, 2014. 44(12): p. 1133–40.

6. Muller, S., et al., Brain Metastases from Colorectal Cancer: A Systematic Review of the Literature and Meta-Analysis to Establish a Guideline for Daily Treatment. Cancers (Basel), 2021. 13(4).

7. Bakhoum, S.F. and L.C. Cantley, The Multifaceted Role of Chromosomal Instability in Cancer and Its Microenvironment. Cell, 2018. 174(6): p. 1347–1360.

8. Klockner, T.C. and C.S. Campbell, Selection forces underlying aneuploidy patterns in cancer. Mol Cell Oncol, 2024. 11(1): p. 2369388.

9. Sdeor, E., et al., Aneuploidy as a driver of human cancer. Nat Genet, 2024. 56(10): p. 2014–2026.

10. Taylor, A.M., et al., Genomic and Functional Approaches to Understanding Cancer Aneuploidy. Cancer Cell, 2018. 33(4): p. 676–689 e3.

11. Brastianos, P.K., et al., Genomic Characterization of Brain Metastases Reveals Branched Evolution and Potential Therapeutic Targets. Cancer Discov, 2015. 5(11): p. 1164–1177.

12. Golas, M.M., et al., Cytogenetic signatures favoring metastatic organotropism in colorectal cancer. Nat Commun, 2025. 16(1): p. 3261.

13. Nguyen, B., et al., Genomic characterization of metastatic patterns from prospective clinical sequencing of 25,000 patients. Cell, 2022. 185(3): p. 563–575 e11.

14. Hu, Z., et al., Quantitative evidence for early metastatic seeding in colorectal cancer. Nat Genet, 2019. 51(7): p. 1113–1122.

15. Zheng, G.X.Y., et al., Haplotyping germline and cancer genomes with high-throughput linked-read sequencing. Nat Biotech, 2016. 34(3): p. 303–311.

16. Li, H. and R. Durbin, Fast and accurate short read alignment with Burrows-Wheeler transform. Bioinformatics, 2009. 25(14): p. 1754–60.

17. McKenna, A., et al., The Genome Analysis Toolkit: A MapReduce framework for analyzing next-generation DNA sequencing data. Genome Research, 2010. 20(9): p. 1297–1303.

18. Casper, J., et al., The UCSC Genome Browser database: 2026 update. Nucleic Acids Res, 2026. 54(D1): p. D1331–D1335.

19. Spurr, L.F., et al., Quantification of aneuploidy in targeted sequencing data using ASCETS. Bioinformatics, 2021. 37(16): p. 2461–2463.

20. Korthauer, K., et al., A practical guide to methods controlling false discoveries in computational biology. Genome Biol, 2019. 20(1): p. 118.

21. Jee, J., et al., Automated real-world data integration improves cancer outcome prediction. Nature, 2024. 636(8043): p. 728–736.

22. Mayakonda, A., et al., Maftools: efficient and comprehensive analysis of somatic variants in cancer. Genome Res, 2018. 28(11): p. 1747–1756.

23. Tagore, S., et al., Single-cell and spatial genomic landscape of non-small cell lung cancer brain metastases. Nat Med, 2025. 31(4): p. 1351–1363.

24. Xing, X., et al., Pan-cancer human brain metastases atlas at single-cell resolution. Cancer Cell, 2025. 43(7): p. 1242–1260 e9.

25. Biermann, J., et al., Dissecting the treatment-naive ecosystem of human melanoma brain metastasis. Cell, 2022. 185(14): p. 2591–2608 e30.

26. Sathe, A., et al., Colorectal cancer relies on an immunosuppressive cellular topography and genomic adaptations for establishing brain metastases. bioRxiv, 2025.

27. Zaalberg, I.C., et al., Aneuploidy patterns in colorectal cancer. Cell Rep, 2026. 45(1): p. 116721.

28. Christensen, T.D., et al., Systematic review: brain metastases from colorectal cancer--Incidence and patient characteristics. BMC Cancer, 2016. 16: p. 260.

29. Li, J., et al., Non-cell-autonomous cancer progression from chromosomal instability. Nature, 2023. 620(7976): p. 1080–1088.

30. Bonadio, R.C., et al., Epidemiology and Outcomes of Patients With Brain Metastases From Colorectal Cancer—Who Are These Patients? Clinical Colorectal Cancer, 2021. 20(3): p. e195–e200.

31. Chang, Y., et al., Survival Outcome of Surgical Resection vs. Radiotherapy in Brain Metastasis From Colorectal Cancer: A Meta-Analysis. Front Med (Lausanne), 2022. 9: p. 768896.

